# Development of Echovirus 29 cytopathology in RD cell line might not happen within 10 days post inoculation

**DOI:** 10.1101/186205

**Authors:** M.O. Adewumi, T.O.C. Faleye, O.T. Ayinde, U.I. Ibok, J.A. Adeniji

**Affiliations:** Department of Virology, College of Medicine, University of Ibadan, Ibadan, Oyo State, Nigeria.; Department of Microbiology, Faculty of Science, Ekiti State University, Ado-Ekiti, Ekiti, State, Nigeria.; WHO National Polio Laboratory, University of Ibadan, Ibadan, OyoState, Nigeria.

**Keywords:** Enterovirus, Echovirus 29, E29, NPEVs, Nigeria

## Abstract

Echovirus 29 (E29) is a member of Species Enterovirus B (EV-B) in the genus Enterovirus, family *Picornaviridae*, order *Picornavirinae*. In Nigeria, molecular characterization of E29 was first described in 2002. In 2015, we found that a new clade of E29 had replaced that described in Nigeria in 2002-2003. To date, E29 isolates described from Nigeria were isolated in cultures of RD cell line. In 2016, we characterised an E29 strain that did not show cytopathology on RD cell line within the recommended 10 days of culture.

Here we show that the E29 in question grows with evident CPE in RD cell culture like other members of the clade when allowed to stay in culture for 13 to 14 days. The findings of this study therefore suggest that some of the samples declared negative for enteroviruses by the current WHO cell culture based detection algorithm might be false negatives. It is therefore encouraged that those particularly interested in non-polio enteroviruses endeavour to maintain at least 14 days incubation in cell culture in a bid to accommodate NPEVs like E29 that might need longer time to develop CPE especially when present at low titre.

## Introduction

Echovirus 29 (E29) is a member of Species Enterovirus B (EV-B) in the genus Enterovirus, family *Picornaviridae*, order *Picornavirinae*. The best known and studied member of the genus is poliovirus and is a member of Species Enterovirus C (EV-C). The E29 virion is a non-enveloped icosahedron with a diameter of 28-30nM. Within the virion is a 5^*l*^-protein-linked, single-stranded, positive sense, RNA genome that is ~7,500 nucleotides long. The genome encodes a single open reading frame (ORF) that is flanked on both ends by untranslated regions (5^*l*^& 3^*l*^UTRs).

Since the 1950s (Rosen et al., 1964), E29s have been described globally and have been recovered from both humans (Oyero et al., 2014) and non-human primates (Sadeuh-Mba et al., 2014). They have been associated with an array of clinical manifestations that include non-specific febrile illnesses (Rosen et al., 1964) and acute flaccid paralysis (Oyero et al., 2014).

In Nigeria, molecular characterization of E29 was first described in 2002 (Oyero et al., 2014) (Figure 1). By 2015, we found (unpublished) that a new clade of E29 had replaced (Figure 1) that found in Nigeria in 2002-2003 (Oyero et al., 2014). In 2017 (unpublished), we found another member of the new clade we first detected in 2015 thereby confirming the presence and circulation of this clade in the region (Figure 1). All the E29 isolates from Nigeria described so far were isolated in cultures of RD cell line (Oyero et al., 2014, Adeniji et al., unpublished).

**Figure 1:**
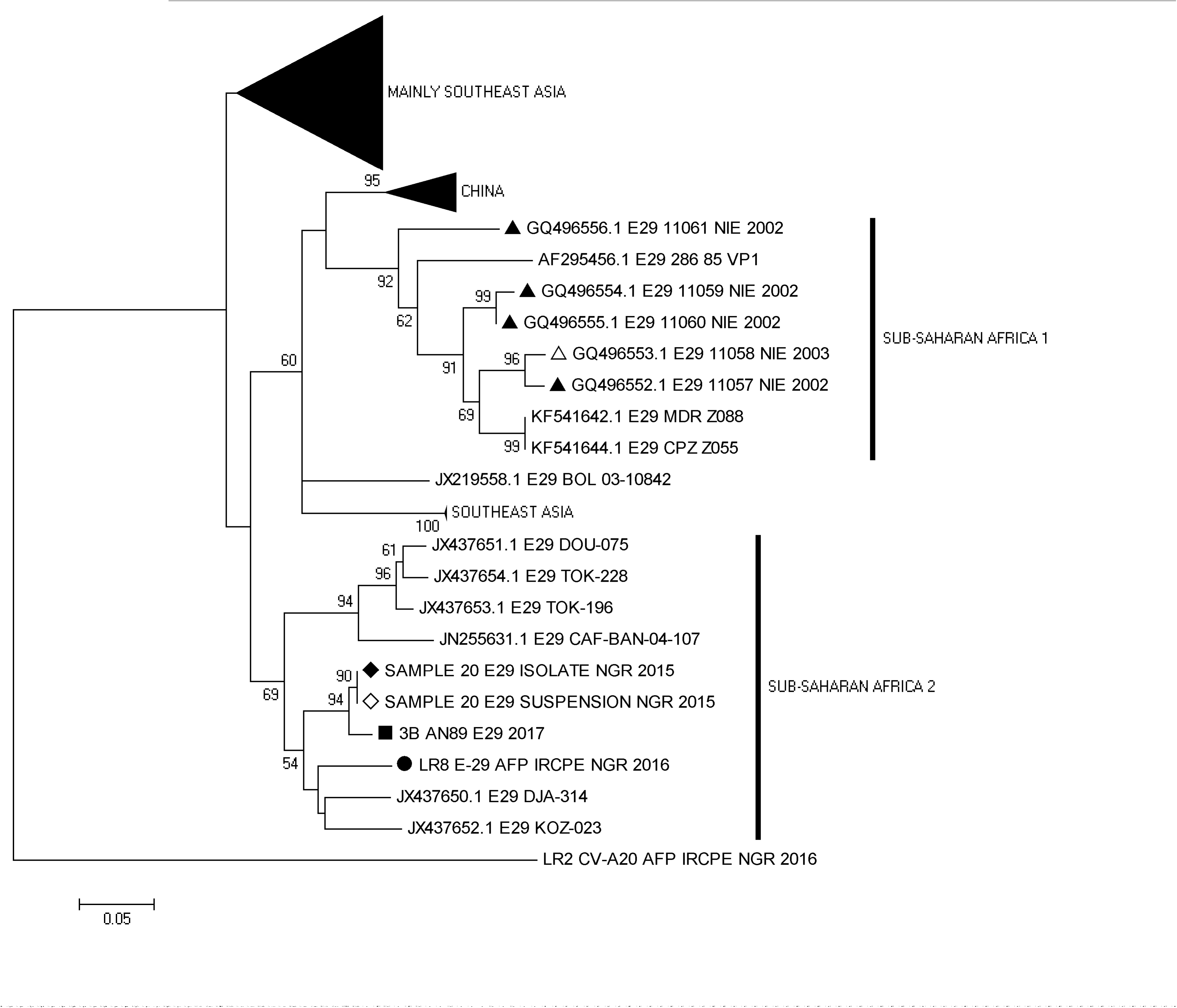
Phylogram of genetic relationship between VP1 nucleotide sequences of E29 isolates. The phylogenetic tree is based on an alignment of the partial VP1 sequences. The strains described are indicated with; black triangle for 2002 Nigerian isolates, white triangle for 2003 Nigerian isolate, black and white diamonds for 2015 Nigerian strains recovered from RD cell culture isolate and the corresponding faecal suspension, respectively, black square for 2017 Nigerian isolate and black circle for the 2016 Nigerian strain discussed in this study. The GenBank accession numbers and strain of the isolates are indicated in the tree. Bootstrap values are indicated if >50%.

Most often, non-polio enteroviruses are isolated on RD (derived from a human rhabdomyosarcoma; McAllister et al. 1969) cell line courtesy the Global Polio Eradication Initiative (GPEI) which has about 150 WHO accredited laboratories globally (Global Polio Laboratory Network [GPLN]) that use the RD cell line for enterovirus isolation as recommended by the WHO (2003, 2004). Enteroviruses show classic cytopathic effect (CPE) in the cell line that includes rounding-up and becoming refractive. The GPEI also uses L20B cell line in its algorithm and particularly for poliovirus isolation. The L20B cell line is a recombinant cell line of mouse origin. It is however engineered to express CD155 (the human poliovirus receptor) (Pipkin et al. 1993).

In 2016 (unpublished), we characterised an E29 strain that did not show cytopathology on RD cell line within the recommended 10 days of culture (http://polioeradication.org/wp-content/uploads/2017/05/NewAlgorithmForPoliovirusIsolationSupplement1.pdf).In fact, this E29 was only detected because it showed CPE on L20B cell line on initial inoculation but the CPE was not reproducible on passage in both L20B and RD cell line. Molecular characterization of the cell culture supernatant of the passage then revealed the presence of E29. This ‘non-reproducible CPE’ is classically associated with Adenoviruses (Thorley and Roberts, 2016), Reoviruses and other non-enteroviruses (http://polioeradication.org/wp-content/uploads/2017/05/NewAlgorithmForPoliovirusIsolationSupplement1.pdf).

What fascinated us was the fact that the E29 strain we found in 2016 belonged to the new clade we recently found to be circulating in the region (figure 1). Why then, unlike other members of the clade, was this E29 strain not showing CPE in RD cell line.

## Methods

To answer this question, we passaged the supernatant (which was previously sequenced to identify the virus) in RD cell line at 37°C for another seven (7) days (Adeniji and Faleye, 2014).The inoculated RD cell culture tubes were microscopically examined everyday for the presence of CPE. After day seven, the tubes were freeze-thawed three times and ten-fold (X10) serial dilutions of the RD cell culture supernatant was made. The stock, 10^−2^, 10^−4^ and 10^−6^ dilutions were then passaged in RD cell line for another 48 hours (Table 1). The inoculated RD cell culture tubes were also microscopically examined every day for the presence of CPE Subsequently, the stock, dilutions and their corresponding supernatant recovered from the 48-hour passaged in RD cell line were freeze thawed three times and stored at −20°C for further analysis.

**Table 1:** Cytopathic effect development after further passage in RD cell line for 48hours and change in virus concentration measured by Real Time RT-PCR.

| S/N | LAB ID | Dilution | Initial Ct Value | Ct Value after passage in RD for 48 hours | Change in Ct Value | CPE in RD cell line (48 hours) |
| --- | --- | --- | --- | --- | --- | --- |
| 1 | LR-8-E-29 | stock | 20.0 | 19.0 | -1.0 | ++++ |
|  |  | 10<sup>-2</sup> | 28.5 | 18.4 | -10.1 | ++++ |
|  |  | 10<sup>-4</sup> | 32.1 | 40.0 | +7.9 | -- |
|  |  | 10<sup>-6</sup> | 35.1 | 40.0 | +4.9 | -- |

RNA was extracted from the stock, dilutions and their corresponding supernatant recovered from the 48-hour passaged in RD cell line, using an RNA extraction kit (JenaBioscience, Jena, Germany). Subsequently, the RNA extract was subjected to the one-step PanEnterovirus real-time Reverse Transcriptase Polymerase Chain Reaction (rRT-PCR) assay currently in use by the GPLN (Zaidi et al., 2016) and interpreted according to the manufacturer’s instructions.

## Result

By day six (6) of passage in RD cell line, CPE started developing in culture tube and by day seven (7), the CPE was 100% (i.e. 4+). Prior this passage, the faecal suspension had been initially inoculated into L20B cell line and incubated at 37°C for 5 days after which it was passaged in RD cell line and incubated at the same temperature for another two days (i.e. it had already been in culture for seven days). Hence, CPE started developing at day 13 of incubationin cell culture and was complete by day 14.

On further passage of the stock, 10^−2^, 10^−4^ and 10^−6^ dilutions in RD cell line for another 48 hours, the stock and 10^−2^ dilution developed complete CPE in the said time but not the 10^−4^ and 10^−6^ dilutions (Table 1). In fact the 10^−4^ and 10^−6^ dilutions had not shown any trace of CPE as at 48 hours when the assay was terminated (Table 1). Result of the rRT-PCR assay showed a drop in Ct value of the stock and 10^−2^ dilution in 48 hours, thereby showing a rise in virus titre and consequently confirming replication. On the other hand, an increase in Ct value was observed for the 10^−4^ and 10^−6^ dilutions, implying a reduction in virus titre and thereby suggesting absence of or very slow replication.

## Discussion

Here we show that the E29 in question grows in RD cell culture with evident CPE like all the other members of the clade (Figure 1). Our findings suggest that the growth dynamics of the virus is very much dependent on its titre and at low titer, the virus might need to stay in culture for longer (13 to 14 days) than the new WHO specified 10 days for CPE to be observed. It is crucial to re-iterate that the 2002-2003 Nigerian isolates were recovered using the WHO cell culture algorithm which require incubation in cell culture for 14 days (WHO, 2003, 2004). Hence even at low titre, there is sufficient time for the virus to accumulate enough variations to adapt to the cell line and replicate efficiently with evident CPE. Against this backdrop, it is likely that the reason the polio laboratory could not detect the E29 in question was because the titre was very low and the new algorithm (http://polioeradication.org/wp-content/uploads/2017/05/NewAlgorithmForPoliovirusIsolationSupplement1.pdf) requires incubation in cell culture for only 10 days which might not be sufficient for development of Echovirus 29 CPE.

The findings of this study therefore suggest that some of the samples declared negative for enteroviruses by the current WHO cell culture based detection algorithm (http://polioeradication.org/wp-content/uploads/2017/05/NewAlgorithmForPoliovirusIsolationSupplement1.pdf) are false negatives. In fact, many of those samples might already be at the brink of developing CPE when their growth in cell culture was terminated and discarded. It is therefore encouraged that those particularly interested in Non-Polio Enteroviruses endeavour to maintain at least 14 days incubation in cell culture in a bid to accommodate NPEVs like E29 that might need 13-14 days to develop CPE especially when present at low titre.

## Acknowledgement

We thank the entire staff of the WHO National Polio Laboratory in Ibadan Nigeria for their valuable technical assistance and RIVM, The Netherlands, for the enterovirus antisera panel.

## Author Contributions

1. StudyDesign (All Authors)

2. SampleCollection, Laboratory and Data analysis (All Authors)

3. Wrote, revised, read and approved the final draft oftheManuscript (All Authors)

